# Molecular correlates of swarming behaviour in *Aedes aegypti* males

**DOI:** 10.1101/2024.05.03.591282

**Authors:** Julien Devilliers, Hollie Marshall, Ben Warren, Charalambos P. Kyriacou, Luciana Ordunha Araripe, Rafaela V. Bruno, Ezio Rosato, Roberto Feuda

**Affiliations:** Neurogenetics Group, University of Leicester, Leicester, UK; Department of Genetics and Genome Biology, College of Life Sciences, University of Leicester, Leicester, United Kingdom; Laboratório de Biologia Molecular de Insetos, Instituto Oswaldo Cruz (Fiocruz), Rio de Janeiro, Brazil; Insitituto Nacional de Ciência e Tecnologia em Entomologia Molecular (INCT-EM)/CNPq, Brazil

**Keywords:** *Aedes aegypti*, swarming, transcriptomics, circadian clock

## Abstract

Mosquitoes are the deadliest vectors of disease. They impose a huge health burden on human populations spreading parasites as disparate as protozoans (malaria), viruses (yellow fever and more), and nematodes (filariasis), which cause life-threatening conditions. In recent years, mating has been proposed as a putative target for population control. Mosquitoes mate mid-air in swarms initiated by males and triggered by a combination of internal and external stimuli. As the number of females in a swarm is limited, there is intense competition among males. Indeed, males ‘retune’ their physiology for this demanding behaviour but the underlying genetic changes are largely unknown. Interestingly, recent evidence indicates that the upregulation of circadian clock genes may be involved in the swarming of malaria mosquitoes of the genus *Anopheles*. Here, we use whole-head RNA-seq to identify gene expression changes in *Aedes aegypti* males who are engaged in swarming in a laboratory setting. Our results suggest that swarming males tend to lower some housekeeping functions while increasing the remodelling of the cytoskeleton and neuronal connectivity; the transcription of circadian clock genes is unaffected.

## INTRODUCTION

Mosquitoes are important insect vectors as they transmit the microorganisms that cause malaria, filariasis, and many viral diseases [1]. In the past century, the main strategies for population control have relied on insecticides and targeted hematophagous females. Currently, widespread resistance to pesticides and increased environmental concerns are urging the adoption of new approaches [2]. One promising strategy advocates using molecular tools to target key genetic regulators of development or reproduction [3,4]. With this approach, genetically modified males (non-biting) are mass-reared and released into the environment, where mating with resident females causes the local population to crash [3]. The underlying principle is that such males would compete effectively with their wild conspecifics to gain multiple access to females, which, depending on the species, tend to resist remating [5,6]. Swarming is the gathering of a large group of males flying together to attract females and competing for copulation. In mosquitoes, the rivalry is particularly fierce. For instance, in *Anopheles gambiae* only 15% *circa* of males form copula pairs [7], while swarming depletes about 50% of their sugar and glycogen reserves [8]. Swarming is initiated by males and is prompted by a combination of internal and external stimuli, such as sex drive, visual landmarks, light levels, time of day, acoustic signals, and olfactory cues [9]. However, the relative importance of these stimuli for swarming is still debated [10–15]. Additionally, we know little about how males ‘tune’ their physiology in preparation for this fundamental behaviour and the genetic regulations behind it. Such knowledge could provide clues for identifying the initial triggers of swarm formation and suggest measures to evaluate the fitness of mass-reared males before release.

We investigated changes in gene expression in swarming males of *Aedes aegypti*, a species with a wide geographical distribution and notable for spreading arboviral diseases, such as dengue, Zika, chikungunya and urban yellow fever [1]. *Aedes aegypti* are diurnal mosquitoes, and swarming can occur at any time of day, although it is more common towards dusk and dawn if a host is unavailable [16]. A swarm forms when tens of males start flying in an erratic figure 8 pattern around a host that provides olfactory cues [17]. Acoustic stimuli are also important, as the acoustic signal generated by a male beating his wings in preparation for swarming may prompt nearby males to join [18]. Males aggregate, and their pheromones attract females to enter the swarm [14]. During flight, females are recognised and chased based on their flight tone, which has a lower frequency than males [18]. Crucially, at the time of swarming, males increase the frequency of their flight tone, reaching ∼1.5 times that of females, which is optimal for identifying them within the swarm [18,19]. This change is partially driven by the circadian clock and is independent of previous interactions with females [20].

We performed mRNA-sequencing (RNA-seq) on whole heads of swarming and non-swarming *Aedes aegypti* males in order to detect transcriptional changes occurring during swarming. We identified 27 differentially expressed transcripts; 11 were upregulated and 16 downregulated in swarming individuals. The changes indicate that swarming males tend to increase the remodelling of the cytoskeleton and neuronal connectivity while reducing some housekeeping processes. Compared to previous work investigating swarming in *Anopheles* [21], we did not identify any overlap between the two species in the differentially expressed genes. Furthermore, in *Aedes aegypti*, the expression of the canonical clock genes was unaffected, although we observed upregulation of *slowpoke*, a gene previously implicated in the synchronisation of circadian rhythmicity among neuronal clusters [22], and of the serotonin receptor *5-HT2A*, involved in the regulation of circadian behaviour [23]. Our results suggest that in *Aedes aegypti* the expression of clock genes does not change at the transcriptional level during swarming, at least in laboratory conditions. Yet, the changes that lead to sustained swarming behaviour possibly involve the clock neuronal network.

## MATERIALS AND METHODS

### Mosquitoes rearing

Animals were reared at the Laboratório de Biologia Molecular de Insetos, Instituto Oswaldo Cruz (Fiocruz, Rio de Janeiro, Brazil), under 12h light – 12h dark (LD 12:12) conditions at 25 °C. Pupae of *Ae. aegypti* (Rockefeller lineage) were provided by the Laboratório de Biologia, Controle e Vigilância de Insetos Vetores, IOC, Fiocruz. Pupae were maintained in plastic cups filled with water inside an experimental cage (metal frame of 40cm x 40cm x 40cm with screens on all sides). After emergence, adults were fed a 10% sucrose solution provided *ad libitum* inside the cage. Most pupae were males, but a few adult females (not sampled) were observed inside the cage and left *in situ* during the experiment.

### Sampling and RNA extraction

We define as swarming males those that aggregated in sustained flight in small groups at the centre of the cage. The proximity of the host was a stimulus for the males to engage in the behaviour. Irresponsive males standing on the cage walls were considered non-swarming. Four independent samples were collected (on the 17^th^, 24^th^ & 31^st^ of May and on the 7^th^ of June 2021), each consisting of 25 swarming and 25 non-swarming males from the same cage. In total, we collected 100 individuals per condition on the 4^th^ day post-emergence, at ZT10.5 – 11 (*Zeitgeber* Time, the time of the *time-giver*, is defined by the light switch, with ZT0 being lights on and ZT12 lights off). This sampling time corresponds to the activity peak in *Aedes aegypti* males [24]. Males were captured by aspiration and snap-frozen in liquid nitrogen. Heads were separated from bodies on ice and stored in TRIzol (Invitrogen) at -80 °C until mRNA extraction. Mosquito heads were homogenised in TRIzol with a motor-operated plastic pestle while thawing. The aqueous phase was isolated with the help of Phasemaker™ tubes (Thermo Fisher Scientific), and the RNA was precipitated with isopropanol using glycogen (Roche) as carrier. A DNA-free kit (Invitrogen) was used to remove any contaminant genomic DNA. Sample quality was assessed using an Agilent RNA 6000 Nano Kit on an Agilent BioAnalyzer 2100 (Agilent Technologies), and a Qubit RNA BR Assay Kit on a Qubit 2.0 Fluorometer (Invitrogen). The eight samples were sequenced by NovoGene (UK) on a NovaSeq 6000 using 150 paired-end technology. The coverage was > 65 million reads/sample.

### Reads mapping, differential gene expression and Gene Ontology enrichment analysis

We used FastQC 0.11.9 [25] to evaluate read quality and MultiQC 1.12 [26] to summarise results. Reads were aligned to the *Aedes aegypti* reference transcriptome (AeaegyptiLVP_AGWG, version 5.7) from VectorBase [27], and their counts were inferred using Kallisto 0.44.0 [28]. We identified differential expression between swarming and non-swarming males with DESeq2 version 1.40.2 [29] implemented in R version 4.3.0 [30]. Transcripts with at least 10 counts were filtered, and samples were normalised by library size with the rlog transformation in DESeq2. We estimated differential transcript expression using a generalised linear model considering data dispersion between samples. We corrected *p-values* according to the Benjamini-Hochberg method [31]. For the differentially expressed genes, a Gene Ontology enrichment analysis was performed with g:Profiler [32], using the expressed genes in the transcriptome as the background. We used VectorBase [27] and OrthoDB [33] to identify orthologs in *Drosophila melanogaster*, and FlyBase release FB2024_01 [34] to find information on gene function.

## RESULTS

We extracted RNA from the whole head, including the antennae, olfactory palps and proboscis, and sensory appendages used for host-seeking, swarming and mating. The mRNA was sequenced to identify genes that may underlie physiological changes sustaining swarming in *Aedes aegypti*. We sampled swarming and non-swarming males toward the end of the day (ZT10.5 – 11.00, in a LD 12:12 regime), a time when locomotor activity peaks and the probability of swarming increases (Figure 1A) [24,35]. We identified 27 differentially expressed transcripts (p<0.05, Log_2_FC>|0.75|); 11 were upregulated and 16 downregulated in swarming individuals (Figure 1B and Table 1). Contrary to previous findings in *Anopheles coluzzii* (part of the *gambiae* complex)[21], the clock genes *period* (*per*) and *timeless* (*tim*) were not upregulated in swarming males (Figure 1B). Differentially expressed genes were annotated using VectorBase [27]. The annotation classified 17 of the 27 transcripts as ‘unspecified products’, of which 5 (1 upregulated and 3 downregulated) were reported as lacking orthologs in *Drosophila melanogaster*. However, using OrthoDB [33] we identified fly orthologs for 3 of them (Table 1). We used the fly orthologs and FlyBase [34] to infer gene function.

**Table 1.**
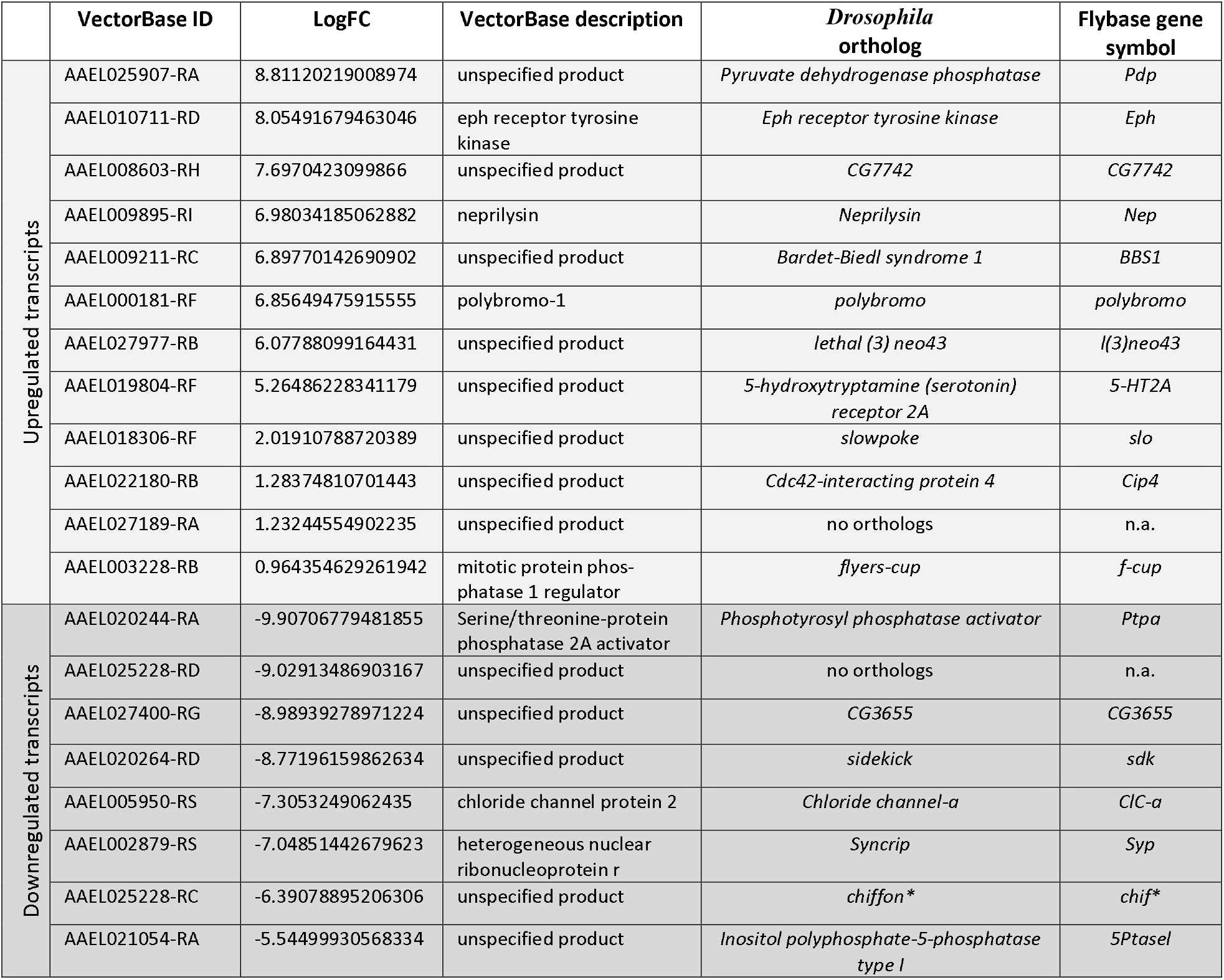
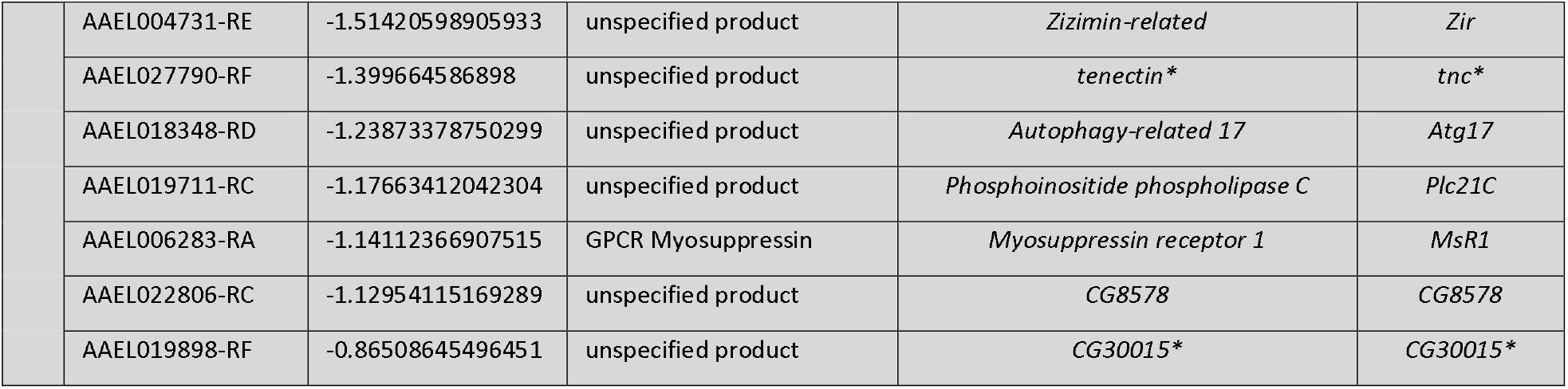
List of genes upregulated and downregulated in swarming *Aedes aegypti* males. *Drosophila* orthologs and their function are also reported. * Indicates orthologs identified using OrthoDB [33].

**Table 2.** List of enriched Gene Ontology (GO) terms linked with upregulated and downregulated transcripts in swarming *Aedes aegypti* males. GO terms for the domains molecular function (MF) and biological process (BP) are shown. The statistical significance of the enrichment is reported in the P_adj_ column. Results are from g:Profiler [32].

| | GO domain | GO term ID | GO term name | $P_{adj}$ |
| --- | --- | --- | --- | --- |
| Upregulated transcripts | BP | GO:0048013 | ephrin receptor signaling pathway | 1,05E-02 |
|  | BP | GO:0006793 | phosphorus metabolic process | 2,81E-02 |
|  | BP | GO:0006796 | phosphate-containing compound metabolic process | 2,81E-02 |
|  | BP | GO:0007169 | transmembrane receptor protein kinase signaling pathway | 2,81E-02 |
|  | BP | GO:0036211 | protein modification process | 2,81E-02 |
|  | BP | GO:1905515 | non-motile cilium assembly | 2,81E-02 |
|  | BP | GO:0006470 | protein dephosphorylation | 2,86E-02 |
|  | BP | GO:0007167 | enzyme-linked receptor protein signaling pathway | 2,86E-02 |
|  | BP | GO:0043412 | macromolecule modification | 2,86E-02 |
|  | BP | GO:0016311 | dephosphorylation | 3,31E-02 |
|  | BP | GO:0019538 | protein metabolic process | 4,62E-02 |
|  | MF | GO:0005003 | ephrin receptor activity | 9,92E-03 |
|  | MF | GO:0060072 | large conductance calcium-activated potassium channel activity | 2,23E-02 |
|  | MF | GO:0004714 | transmembrane receptor protein tyrosine kinase activity | 2,48E-02 |
|  | MF | GO:0004722 | protein serine/threonine phosphatase activity | 2,48E-02 |
|  | MF | GO:0005227 | calcium-activated cation channel activity | 2,48E-02 |
|  | MF | GO:0015269 | calcium-activated potassium channel activity | 2,48E-02 |
|  | MF | GO:0019199 | transmembrane receptor protein kinase activity | 2,48E-02 |
|  | MF | GO:0140096 | catalytic activity, acting on a protein | 2,48E-02 |
|  | MF | GO:0004721 | phosphoprotein phosphatase activity | 3,24E-02 |
|  | MF | GO:0004713 | protein tyrosine kinase activity | 3,24E-02 |
|  | MF | GO:0003682 | chromatin binding | 3,24E-02 |
| Downregulated transcripts | BP | GO:0000422 | autophagy of mitochondrion | 2,50E-02 |
|  | BP | GO:1903008 | organelle disassembly | 2,50E-02 |
|  | BP | GO:0061726 | obsolete mitochondrion disassembly | 2,50E-02 |
|  | MF | GO:0052743 | inositol tetrakisphosphate phosphatase activity | 5,87E-03 |
|  | MF | GO:0004445 | inositol-polyphosphate 5-phosphatase activity | 5,87E-03 |
|  | MF | GO:0052659 | inositol-1,3,4,5-tetrakisphosphate 5-phosphatase activity | 5,87E-03 |
|  | MF | GO:0052658 | inositol-1,4,5-trisphosphate 5-phosphatase activity | 5,87E-03 |
|  | MF | GO:0046030 | inositol trisphosphate phosphatase activity | 5,87E-03 |
|  | MF | GO:0019211 | phosphatase activator activity | 5,87E-03 |
|  | MF | GO:0019208 | phosphatase regulator activity | 9,34E-03 |
|  | MF | GO:0005247 | voltage-gated chloride channel activity | 1,26E-02 |
|  | MF | GO:0003755 | peptidyl-prolyl cis-trans isomerase activity | 1,37E-02 |
|  | MF | GO:0016859 | cis-trans isomerase activity | 1,37E-02 |
|  | MF | GO:0008308 | voltage-gated monoatomic anion channel activity | 1,37E-02 |
|  | MF | GO:0052745 | inositol phosphate phosphatase activity | 1,58E-02 |
|  | MF | GO:0004435 | phosphatidylinositol phospholipase C activity | 1,58E-02 |
|  | MF | GO:0004629 | phospholipase C activity | 1,58E-02 |
|  | MF | GO:0008047 | enzyme activator activity | 2,01E-02 |
|  | MF | GO:0042578 | phosphoric ester hydrolase activity | 2,28E-02 |
|  | MF | GO:0016853 | isomerase activity | 2,28E-02 |
|  | MF | GO:0005254 | chloride channel activity | 2,28E-02 |
|  | MF | GO:0030234 | enzyme regulator activity | 2,28E-02 |
|  | MF | GO:0005253 | monoatomic anion channel activity | 2,38E-02 |
|  | MF | GO:0015108 | chloride transmembrane transporter activity | 2,62E-02 |
|  | MF | GO:0140677 | molecular function activator activity | 2,62E-02 |
|  | MF | GO:0098772 | molecular function regulator activity | 2,94E-02 |
|  | MF | GO:0015103 | inorganic anion transmembrane transporter activity | 3,03E-02 |
|  | MF | GO:0022832 | voltage-gated channel activity | 3,37E-02 |
|  | MF | GO:0005244 | voltage-gated monoatomic ion channel activity | 3,37E-02 |
|  | MF | GO:0008509 | monoatomic anion transmembrane transporter activity | 3,61E-02 |

**Figure 1.**
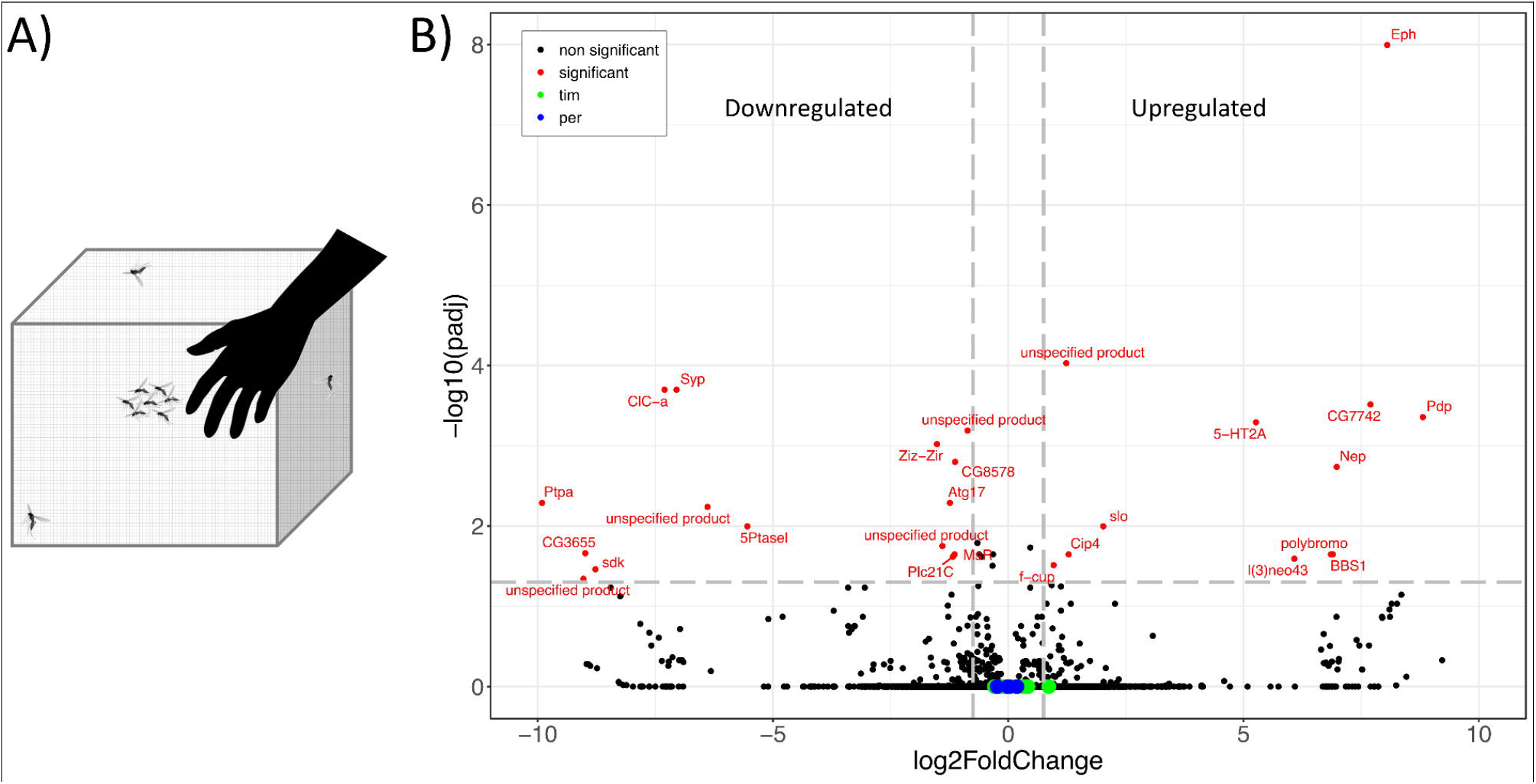
A) Experimental design. Swarming males (flying) are attracted to a human host. Non-swarming males are irresponsive to the host and rest on the walls of the cage. B) Volcano plot showing transcripts differentially expressed in swarming males (in red) (p<0.05, Log2FC>|0.75|). Transcripts of the circadian genes *period* (*per*) and *timeless* (*tim*) and are not differentially expressed and are shown in blue and green, respectively.

In swarming males, upregulated genes included *Pyruvate dehydrogenase phosphatase* (*Pdp*, AAEL025907-RA) that is involved in both metabolic and signalling pathways, *Ephrin tyrosin kinase receptor* (*Eph*, AAEL010711-RD) whose product promotes cell adhesion and axon guidance, *polybromo* (AAEL000181-RF) encoding a chromatin remodeler, and *serotonin receptor 2A* (*5-HT2A*, AAEL019804-RF) implicated in food choice, ageing and modulation of circadian behaviour [23,36]. Additional genes of interest were *slowpoke* (*slo*, AAEL018306-RF), whose product is a potassium ion channel (BK type) involved in odour recognition in *Anopheles gambiae* [37] and in the synchronisation of circadian neurons in flies [22], and *Bardet-Bield syndrome 1* (*BBS1*, AAEL009211-RC) that encodes a protein expressed in sensory neurons in *Drosophila*, important for cilia biogenesis [38]. Indeed, gene ontology analysis revealed an enrichment in the categories of cell communication, ephrin receptor signal pathway and cilium assembly (Tables 1 & 2).

Among the downregulated genes, a common theme was involvement in housekeeping functions. *Phosphotyrosyl phosphatase activator* (*Ptpa*, AAEL020244-RA) is implicated in cell fate determination, mitotic spindle organisation, and basal protein localisation. The gene *sidekick* (SDK, AAEL020264-RD) regulates the remodelling of epithelia and synapses. *Syncrip* (*Syp*, AAEL002879-RS) encodes an RNA-binding protein involved in many biological processes, including the negative regulation of synaptic assembly. The product of *chiffon* (*chif*, AAEL025228-RC) mediates cell division and cell cycle progression. Additional biological processes that may be affected are immune responses (*Zir*, AAEL004731-RE), autophagy (*Atg17*, AAEL018348-RD), contractility of visceral muscles (*MsR1*, AAEL006283-RA), and inositol metabolism (*5Ptasel*, AAEL021054-RA) that is important for energy homeostasis [39]. Accordingly, organelle disassembly, autophagy of mitochondrion (Figure 2C and D), and inositol triphosphate phosphatase activity were enriched ontology terms (Tables 1 & 2).

## DISCUSSION

Swarming is fundamental for the biology of mosquitoes. It marks the time and place where males aggregate to attract females and compete for mating. It is a challenging behaviour involving several sensory modalities and rapid sensory-motor integration; among many males, only the fastest in recognising and catching a female in mid-flight will be able to mate. In this study, we aimed to identify gene expression changes linked with swarming behaviour. Our findings indicate that males involved in swarming undergo cytoskeletal reorganisation, heightened cellular connectivity, and a decrease in various housekeeping functions. These changes are likely facilitated by the activation of chromatin remodelling pathways (Figure 1B and Table 1). It is revealing that swarming in *Aedes aegypti* may require promoting processes expected to redefine cellular performance and connectivity, and we speculate that they contribute to ‘upscaling’ motor abilities and sensory acuity required for swarming. Physiological and structural changes correlated to swarming have been documented for flying and hearing [20,40,41]. These appear to occur for other sensory modalities, such as vision [42].

Recent work carried out in the malaria mosquito *Anopheles coluzzii* identified increased expression of the canonical clock genes *per* and *tim* in swarming males in the field [21]. Additionally, the authors described increased expression for carbohydrate, amino acid, or lipid metabolism genes and an overall reduction for cytoskeletal and structural genes [21]. In *Aedes aegypti*, we did not detect differential expression for clock genes (Figure 1B), and the expression of metabolic and cytoskeletal genes showed an opposite trend (Table 1). Furthermore, we observed no overlap in differentially expressed genes between the two species. We suspect there are important biological reasons that justify these discrepancies although technical considerations (microarrays *versus* RNAseq) may contribute to the differences.

Wang and co-workers [21] collected *Anopheles* mosquitoes at dusk from the wild, sampling males that were swarming outdoors and males that were at rest in houses, two very different environments. We sampled *Aedes* in the laboratory, collecting swarming and non-swarming males from the same cage. Thus, the disparity in the list of genes showing differential expression between *Anopheles* and *Aedes* may reflect environmental rather than, or in addition to, species-specific differences.

Differences between species also contribute. In nocturnal *Anopheles* mosquitoes, swarming is strictly crepuscular. In the laboratory it is maximal around ZT14-15, close to the peaks of *per* and *tim* expression, suggesting a tight regulation by the clock [20,21]. In the diurnal *Aedes aegypti*, the timing of swarming is less strict, although it occurs mainly toward the end of the day, reaching a maximum before dusk (ZT11 in the laboratory)[16]. Thus, in *Aedes aegypti*, we cannot rule out that low transcription of *per* and *tim* at ZT11 could mask changes in their expression linked with swarming.

In our study, we did not detect differential expression of canonical clock genes during swarming. However, concluding that the clock is not involved in swarming may be premature as both *Anopheles* and *Aedes* show a reduction in mating and fertility when the clock is disrupted [21,35], indicating that many processes are affected. Additionally, it is notable that *slowpoke* (*slo*) was upregulated in swarming males. This gene encodes a calcium-dependent potassium channel that regulates circadian locomotor activity in *Drosophila melanogaster* [22,43,44]. Indeed, several groups of (dorsal) circadian neurons are desynchronised in *slo*-null flies due to reduced release of the neuropeptide Pigment Dispersing Factor (PDF). PDF is produced by an important ‘pacemaker’ cluster of clock neurons and functions as a synchronisation agent across the neuronal clock network [44]. Moreover, in flies, SLO is directly involved in acquiring circadian information in neuronal circuits downstream of the clock that are necessary for the maintenance of locomotor activity rhythms [43]. Interestingly, mutations in *slowpoke* cause changes to the courtship song in *Drosophila*, a sound generated by males vibrating their wings [45]. The organisation of the circadian network in *Anopheles coluzzii* and in *Aedes aegypti* is based on the expression of PER and PDF, as with Drosophila, although species-specific differences exist [46]. A further similarity lies in the fact that although they do not generate a song, males of many mosquito species change the frequency of their wingbeat for mating [20]. Thus, it is tempting to speculate that *slo*, like in the fly, may have circadian and ‘mating’ roles in mosquitoes. We suggest that in mosquitoes, the circadian neuronal network supports swarming beyond the direct involvement of the molecular cogs of the clock. Testing this hypothesis will motivate our future work.

## Acknowledgements

This work was funded by grant E-26/201.334/2016 (FAPERJ – Brazil) to RB and by a UKRI Global Challenges Research Fund (M38PF27) to ER, CPK and RF. RF acknowledges a Royal Society University Research Fellowship (UF160226 and URF\R\221011). JD is supported by a PhD scholarship from the College of Life Science (University of Leicester).

## Author contributions

ER, CPK, BW, and RF conceived the study. JD, LOA, and RVB carried out the experiments. JD and HM analysed the data. JD, RF, RVB, and ER wrote the draft manuscript. All authors contributed to the final version.

## Data Accessibility

Data has been deposited in GenBank under NCBI BioProject: XXX. All codes are available at: https://github.com/juliendevi/Swarming_vs_non-swarming_Aedes_aegypti.

## REFERENCES

1. WHO. 2020 Vector-borne diseases.

2. Diabate A, Tripet F. 2015 Targeting male mosquito mating behaviour for malaria control. Parasites Vectors 8, 347. (doi:10.1186/s13071-015-0961-8)

3. Alphey L. 2014 Genetic Control of Mosquitoes. Annu. Rev. Entomol. 59, 205–224. (doi:10.1146/annurev-ento-011613-162002)

4. Hammond A et al. 2016 A CRISPR-Cas9 gene drive system targeting female reproduction in the malaria mosquito vector Anopheles gambiae. Nat Biotechnol 34, 78–83. (doi:10.1038/nbt.3439)

5. Gabrieli P, Kakani EG, Mitchell SN, Mameli E, Want EJ, Mariezcurrena Anton A, Serrao A, Baldini F, Catteruccia F. 2014 Sexual transfer of the steroid hormone 20E induces the postmating switch in Anopheles gambiae. Proc. Natl. Acad. Sci. U.S.A. 111, 16353–16358. (doi:10.1073/pnas.1410488111)

6. Oliva CF, Damiens D, Vreysen MJB, Lemperière G, Gilles J. 2013 Reproductive Strategies of Aedes albopictus (Diptera: Culicidae) and Implications for the Sterile Insect Technique. PLoS ONE 8, e78884. (doi:10.1371/journal.pone.0078884)

7. Sawadogo PS et al. 2014 Swarming behaviour in natural populations of Anopheles gambiae and An. coluzzii: Review of 4 years survey in rural areas of sympatry, Burkina Faso (West Africa). Acta Tropica 132, S42–S52. (doi:10.1016/j.actatropica.2013.12.011)

8. Maïga H, Dabiré RK, Lehmann T, Tripet F, Diabaté A. 2012 Variation in energy reserves and role of body size in the mating system of Anopheles gambiae. Journal of Vector Ecology 37, 289–297. (doi:10.1111/j.1948-7134.2012.00230.x)

9. Clements AN. 1999 The biology of mosquitoes. Volume 2: sensory reception and behaviour.

10. Göpfert MC, Briegel H, Robert D. 1999 Mosquito hearing: sound-induced antennal vibrations in male and female Aedes aegypti. Journal of Experimental Biology 202, 2727–2738. (doi:10.1242/jeb.202.20.2727)

11. Göpfert MC, Robert D. 2000 Nanometre–range acoustic sensitivity in male and female mosquitoes. Proc. R. Soc. Lond. B 267, 453–457. (doi:10.1098/rspb.2000.1021)

12. Gibson G, Russell I. 2006 Flying in Tune: Sexual Recognition in Mosquitoes. Current Biology 16, 1311– 1316. (doi:10.1016/j.cub.2006.05.053)

13. Howell PI, Knols BG. 2009 Male mating biology. Malar J 8, S8. (doi:10.1186/1475-2875-8-S2-S8)

14. Fawaz EY, Allan SA, Bernier UR, Obenauer PJ, Diclaro JW. 2014 Swarming mechanisms in the yellow fever mosquito: aggregation pheromones are involved in the mating behavior of Aedes aegypti. Journal of Vector Ecology 39, 347–354. (doi:10.1111/jvec.12110)

15. Baeshen R. 2022 Swarming Behavior in Anopheles gambiae (sensu lato): Current Knowledge and Future Outlook. Journal of Medical Entomology 59, 56–66. (doi:10.1093/jme/tjab157)

16. Cabrera M, Jaffe K. 2007 An aggregation pheromone modulates lekking behaviour in the vector mosquito Aedes aegypti (Diptera: Culicidae). Journal of the American Mosquito Control Association 23, 1– 10. (doi:10.2987/8756-971X(2007)23[1:AAPMLB]2.0.CO;2)

17. Hartberg WK. 1971 Observations on the Mating Behaviour of Aedes aegypti in Nature. Bull World Health Organ 45, 847–850.

18. Cator LJ, Arthur BJ, Harrington LC, Hoy RR. 2009 Harmonic Convergence in the Love Songs of the Dengue Vector Mosquito. Science 323, 1077–1079. (doi:10.1126/science.1166541)

19. Gesto JSM, Araki AS, Caragata EP, De Oliveira CD, Martins AJ, Bruno RV, Moreira LA. 2018 In tune with nature: Wolbachia does not prevent pre-copula acoustic communication in Aedes aegypti. Parasites Vectors 11, 109. (doi:10.1186/s13071-018-2695-x)

20. Somers J et al. 2022 Hitting the right note at the right time: Circadian control of audibility in Anopheles mosquito mating swarms is mediated by flight tones. Sci. Adv. 8, eabl4844. (doi:10.1126/sciadv.abl4844)

21. Wang G et al. 2021 Clock genes and environmental cues coordinate Anopheles pheromone synthesis, swarming, and mating. Science 371, pp.411–415.

22. Ceriani MF, Hogenesch JB, Yanovsky M, Panda S, Straume M, Kay SA. 2002 Genome-Wide Expression Analysis in Drosophila Reveals Genes Controlling Circadian Behavior. J. Neurosci. 22, 9305–9319. (doi:10.1523/JNEUROSCI.22-21-09305.2002)

23. Nichols CD. 2007 5-HT 2 receptors in Drosophila are expressed in the brain and modulate aspects of circadian behaviors. Developmental Neurobiology 67, 752–763. (doi:10.1002/dneu.20370)

24. Araripe LO, Bezerra JRA, Rivas GB da S, Bruno RV. 2018 Locomotor activity in males of Aedes aegypti can shift in response to females’ presence. Parasites Vectors 11, 254. (doi:10.1186/s13071-018-2635-9)

25. Andrews S. 2010 FastQC: A Quality Control Tool for High Throughput Sequence Data.

26. Ewels P, Magnusson M, Lundin S, Käller M. 2016 MultiQC: summarize analysis results for multiple tools and samples in a single report. Bioinformatics 32, 3047–3048. (doi:10.1093/bioinformatics/btw354)

27. Giraldo-Calderón GI et al. 2015 VectorBase: an updated bioinformatics resource for invertebrate vectors and other organisms related with human diseases. Nucleic Acids Research 43, D707–D713. (doi:10.1093/nar/gku1117)

28. Bray NL, Pimentel H, Melsted P, Pachter L. 2016 Near-optimal probabilistic RNA-seq quantification. Nat Biotechnol 34, 525–527. (doi:10.1038/nbt.3519)

29. Love MI, Huber W, Anders S. 2014 Moderated estimation of fold change and dispersion for RNA-seq data with DESeq2. Genome Biol 15, 550. (doi:10.1186/s13059-014-0550-8)

30. R Core Team. 2024 R: A Language and Environment for Statistical Computing. Vienna, Austria: R Foundation for Statistical Computing. See https://www.R-project.org/.

31. Benjamini Y, Hochberg Y. 1995 Controlling the False Discovery Rate: A Practical and Powerful Approach to Multiple Testing. Journal of the Royal Statistical Society: Series B (Methodological) 57, 289– 300. (doi:10.1111/j.2517-6161.1995.tb02031.x)

32. Kolberg L, Raudvere U, Kuzmin I, Adler P, Vilo J, Peterson H. 2023 g:Profiler—interoperable web service for functional enrichment analysis and gene identifier mapping (2023 update). Nucleic Acids Research 51, W207–W212. (doi:10.1093/nar/gkad347)

33. Zdobnov EM, Kuznetsov D, Tegenfeldt F, Manni M, Berkeley M, Kriventseva EV. 2021 OrthoDB in 2020: evolutionary and functional annotations of orthologs. Nucleic Acids Research 49, D389–D393. (doi:10.1093/nar/gkaa1009)

34. Gramates LS et al. 2022 FlyBase: a guided tour of highlighted features. Genetics 220, iyac035. (doi:10.1093/genetics/iyac035)

35. Shetty V, Meyers JI, Zhang Y, Merlin C, Slotman MA. 2022 Impact of disabled circadian clock on yellow fever mosquito Aedes aegypti fitness and behaviors. Sci Rep 12, 6899. (doi:10.1038/s41598-022-10825-5)

36. Munneke AS, Chakraborty TS, Porter SS, Gendron CM, Pletcher SD. 2022 The serotonin receptor 5-HT2A modulates lifespan and protein feeding in Drosophila melanogaster. Front. Aging 3, 1068455. (doi:10.3389/fragi.2022.1068455)

37. Kwon Y, Kim SH, Ronderos DS, Lee Y, Akitake B, Woodward OM, Guggino WB, Smith DP, Montell C. 2010 Drosophila TRPA1 Channel Is Required to Avoid the Naturally Occurring Insect Repellent Citronellal. Current Biology 20, 1672–1678. (doi:10.1016/j.cub.2010.08.016)

38. Avidor-Reiss T, Maer AM, Koundakjian E, Polyanovsky A, Keil T, Subramaniam S, Zuker CS. 2004 Decoding Cilia Function: Defining Specialized Genes Required for Compartmentalized Cilia Biogenesis. Cell 117, 527–539. (doi:10.1016/s0092-8674(04)00412-x)

39. Chatree S, Thongmaen N, Tantivejkul K, Sitticharoon C, Vucenik I. 2020 Role of Inositols and Inositol Phosphates in Energy Metabolism. Molecules 25, 5079. (doi:10.3390/molecules25215079)

40. Georgiades M et al. 2023 Hearing of malaria mosquitoes is modulated by a beta-adrenergic-like octo-pamine receptor which serves as insecticide target. Nat Commun 14, 4338. (doi:10.1038/s41467-023-40029-y)

41. Su MP, Andrés M, Boyd-Gibbins N, Somers J, Albert JT. 2018 Sex and species specific hearing mechanisms in mosquito flagellar ears. Nat Commun 9, 3911. (doi:10.1038/s41467-018-06388-7)

42. Feuda R et al. 2021 Phylogenomics of Opsin Genes in Diptera Reveals Lineage-Specific Events and Contrasting Evolutionary Dynamics in Anopheles and Drosophila. Genome Biology and Evolution 13, evab170. (doi:10.1093/gbe/evab170)

43. Ruiz D, Bajwa ST, Vanani N, Bajwa TA, Cavanaugh DJ. 2021 Slowpoke functions in circadian output cells to regulate rest:activity rhythms. PLoS ONE 16, e0249215. (doi:10.1371/journal.pone.0249215)

44. Fernández MDLP, Chu J, Villella A, Atkinson N, Kay SA, Ceriani MF. 2007 Impaired clock output by altered connectivity in the circadian network. Proc. Natl. Acad. Sci. U.S.A. 104, 5650–5655. (doi:10.1073/pnas.0608260104)

45. Peixoto AA, Hall JC. 1998 Analysis of Temperature-Sensitive Mutants Reveals New Genes Involved in the Courtship Song of Drosophila. Genetics 148, 827–838. (doi:10.1093/genetics/148.2.827)

46. Baik LS, Nave C, Au DD, Guda T, Chevez JA, Ray A, Holmes TC. 2020 Circadian Regulation of Light-Evoked Attraction and Avoidance Behaviors in Daytime-versus Nighttime-Biting Mosquitoes. Current Biology 30, 3252-3259.e3. (doi:10.1016/j.cub.2020.06.010)

